# CLIQ-BID: A method to quantify bacteria-induced damage to eukaryotic cells by automated live-imaging of bright nuclei

**DOI:** 10.1101/236091

**Authors:** Yann Wallez, Stéphanie Bouillot, Emmanuelle Soleilhac, Philippe Huber, Ina Attrée, Eric Faudry

## Abstract

Pathogenic bacteria induce eukaryotic cell damage which range from discrete modifications of signalling pathways, to morphological alterations and even to cell death. Accurate quantitative detection of these events is necessary for studying host-pathogen interactions and for developing strategies to protect host organisms from bacterial infections. Investigation of morphological changes is cumbersome and not adapted to high-throughput and kinetics measurements. Here, we describe a simple and cost-effective method based on automated analysis of live cells with stained nuclei, which allows real-time quantification of bacteria-induced eukaryotic cell damage at single-cell resolution. We demonstrate that this automated high-throughput microscopy approach permits screening of libraries composed of interference-RNA, bacterial strains, antibodies and chemical compounds in *ex vivo* infection settings. The use of fluorescently-labelled bacteria enables the concomitant detection of changes in bacterial growth. Using this method named CLIQ-BID (Cell Live Imaging Quantification of Bacteria Induced Damage), we were able to distinguish the virulence profiles of different pathogenic bacterial species and clinical strains.

## INTRODUCTION

Bacterial toxins targeting eukaryotic cells can either directly affect plasma membrane integrity or alternatively they may be internalized, translocated or injected inside the cells. Independent of their route, toxins induce modifications of cell morphology and/or provoke host-cell death. For example, the Anthrax Lethal Toxin (LT) is able to provoke pyroptosis or apoptosis, depending on the cell type and the LT concentration. Furthermore, at sub-lethal concentrations, it induces modification of the cytoskeleton and alters the distribution of junction proteins in endothelial and epithelial cells^1^. In Gram-negative bacteria, Type Three Secretion System (T3SS) toxins hijack eukaryotic signalling pathways, leading to damage ranging from modifications of the normal cytoskeleton function, to cell death, depending on the cell type and the toxin^2^.

Host-pathogen interaction studies therefore rely on detection and quantification of the bacteria-induced eukaryotic cell injuries. Plasma membrane permeabilization leading to cell death, the most dramatic outcome of the cell intoxication process, is usually monitored through the enzymatic measurement of lactate dehydrogenase released after plasma membrane rupture, or through the detection of nuclear stain incorporation by flow cytometry^3–5^. However, the analysis of early events such as the morphological changes induced by cytoskeleton rearrangements are usually based on fixed and stained cells, rendering fine kinetics studies laborious, or on expression of fluorescent chimeric markers, a time-consuming procedure to which some cells are refractory^6^. These approaches are not easily accessible to non-expert scientists.

Overall, there is a dearth of simple methods allowing real-time quantification of morphological changes or cell death. Here, we present the CLIQ-BID method, based on automated high-throughput monitoring of the fluorescence intensity of eukaryotic cell nuclei stained with vital-Hoechst. This live-imaging method permits real-time quantification of bacteria-induced cell damage at single-cell resolution. Starting from an observation in the context of the *Pseudomonas aeruginosa* T3SS, it was extended to other Gram-positive and Gram-negative bacteria equipped with diverse virulence factors. Towards identification of new antibacterial therapeutic targets or research tools, this convenient approach could be employed in functional high-throughput screening of interference-RNA, bacterial strains, antibodies or small molecules. More generally, the CLIQ-BID method could also be used in other cytotoxicity and cell-stress studies.

## RESULTS

### *P. aeruginosa* induces a quantifiable nuclei size reduction

The injection of the exotoxins ExoS, T, Y and ExoU by the T3SS machinery is one of the main virulence determinants of *P. aeruginosa* clinical strains^7^. Those toxins have profound effects on eukaryotic cell biology, provoking plasma membrane disruption or inhibition of phagocytosis followed by a delayed apoptosis^8^. Visually, ExoS and ExoT action on host cytoskeleton leads to a reduction of cell area and a “shrinkage” phenotype^9^. In the search for robust descriptors of this phenomenon, we observed that the Hoechst-stained nuclei of Human Umbilical Vascular Endothelial Cells (HUVECs) become gradually smaller and brighter during incubation with the wild-type *P. aeruginosa* strain PAO1 harbouring ExoS and ExoT. In addition this increased intensity of nuclear staining remarkably correlated with the decrease of cell area (Fig. 1a, compare upper and lower images). The built-in Arrayscan image analysis workflow was employed in order to obtain the nuclei mask (Fig. 1a insert – magenta outlines) by intensity thresholding and the quantitative features corresponding to their areas and fluorescence intensities. The graphical representation of these features extracted from 70 nuclei at different time points clearly shows a negative correlation between nuclei area and intensity (Fig. 1b). Indeed, the condensation of the nuclei results in an increased concentration of the fluorescent dye complexed to the DNA and thus in an enhanced fluorescence intensity. Furthermore, a nuclear intensity threshold could readily be set to segregate cells with bright nuclei (Fig. 1c). Therefore, a subpopulation of cells displaying bright nuclei, which corresponds to the shrunk cells, could be automatically identified by monitoring the nuclear staining intensity.

**Figure 1:**
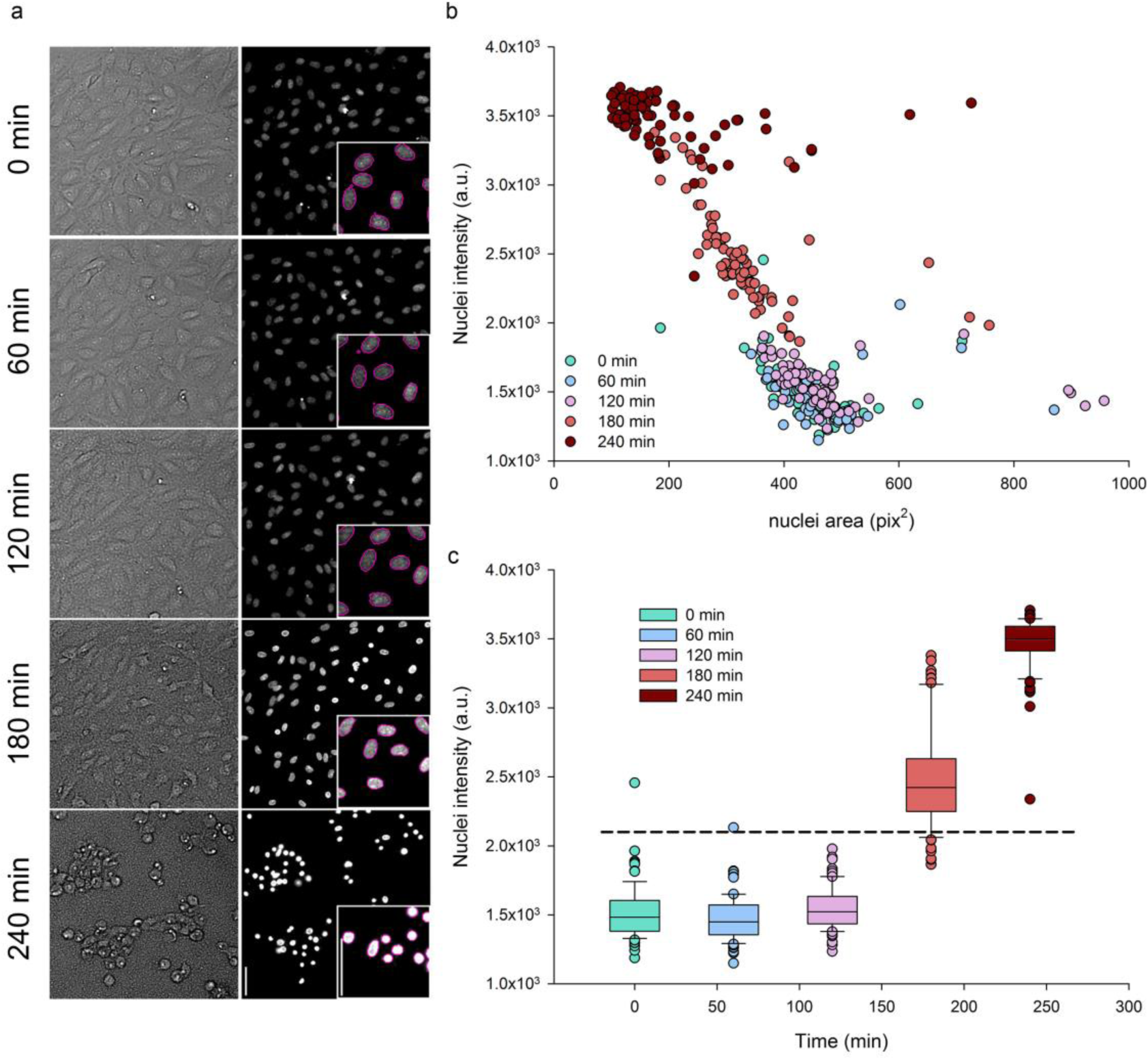
Smaller and brighter cell nuclei reflect *P. aeruginosa* induced cell-damage. Human primary endothelial cells (HUVECs) were infected with *P. aeruginosa* and monitored at different stages of infection by live-imaging microscopy with vital-Hoechst nuclear stain. a) Cell surface and cell nuclei observed in transmitted light and by fluorescent labelling, upper and lower images respectively, in the same field at one hour intervals. Nuclei from the same set of images were automatically segmented (insert – magenta outlines). The scale bars shown on the last nuclei image correspond to 50 μm. b) The area and fluorescence intensities of each segmented nucleus were plotted. Data obtained from different time points are represented in colors, from green (0 min) to dark red (240 min). a.u. = arbitrary units. c) Nuclear staining intensities of cells at different time points of infection are represented in box plots. Whiskers indicate the 10th and 90th percentiles; the top and bottom lines represent the 25th and 75th percentiles; the middle line and dots respectively show median and outliers. Intensities from the three first time points are statistically different from those of the two last time points (one way ANOVA, P < 0.05). The horizontal dashed line represent a threshold that could discriminate between normal and bright nuclei. In b) and c), n = 70 cells at each time point.

In order to determine whether the observed nuclei condensation was a phenomenon restricted to HUVECs, the experiment was repeated on CHO, NIH 3T3, HeLa and A549 cells. Despite slight differences in kinetics between the cell types, the nuclear staining intensity increased during the time course of infection by *P. aeruginosa* (Supplementary Fig. S1 and Supplementary Video S1). Of importance, increased brightness was strictly correlated with cell shrinkage (compare upper and lower images of Supplementary Fig. S1).

*P. aeruginosa* induces a nuclear condensation that is responsible for the observed increase in nuclear staining intensity. A statistical analysis was performed in order to determine which of the nuclear features (i.e. area versus average intensity) better distinguishes between condensed and non-altered populations. To that end, images from uninfected and 3h-infected cells were obtained at 20x and 5x magnifications. Next, the nuclei images were analysed and three features were compared: the mean nuclear area, the mean nuclear intensity and the percentage of bright nuclei, which is the proportion of nuclei whose intensity is above a fixed threshold. These three parameters were the most promising among the different features calculated by the built-in software from the experiment described in Figure 1. The ability of these three parameters to discriminate between the images obtained at the beginning and at the end of the infection was assessed through the Z’-factor (abbreviated Z’). This statistical coefficient takes into consideration the standard deviations as well as the difference between the means of the positive and negative controls, and is often used for the optimization and validation of High Throughput Screening assays^10^. The values reported in Supplementary Table S1 clearly indicate that the percentage of bright nuclei is the most discriminant parameter with Z’=0.64 for 20x magnification, and that taking images at 5x magnification further increases the power of the test (Z’=0.75).

The vital nuclear staining used here offers the possibility to monitor modifications of the nuclei by live-imaging during infection. To obtain live kinetics, the nuclei of the five cell types were labelled prior to infection with *P. aeruginosa* and images were taken at 5x magnification every 15 minutes for 4 hours. Images of the same fields at one hour interval are presented (Fig. 2a). Afterward, automated segmentation of the nuclei and measurement of fluorescence intensities were performed as described above and the percentages of bright nuclei were extracted from the built-in software. To visualize the cells that were considered to be damaged upon bacterial infection, nuclei with intensities above a fixed threshold were delineated in green while those below the threshold were delineated in magenta (Fig. 2a - inserts). Nuclei segmentation and thresholding based on vital-Hoechst staining intensity properly reflected the observed appearance of bright nuclei during the progression of infection. Furthermore, images show that nuclei condensation of 3T3, HeLa and A549 cells occurred earlier than for HUVEC and CHO cells. This was confirmed by the kinetic plots (Fig. 2b) that successfully detected an increase of bright nuclei, occurring exponentially with the duration of cell infection. These plots further depicted differences between cells in terms of inflection time and curve steepness. It was therefore possible to identify kinetics signatures for each of the five cell types. The method was named CLIQ-BID, standing for Cell Live Imaging Quantification of Bacteria Induced Damage.

**Figure 2:**
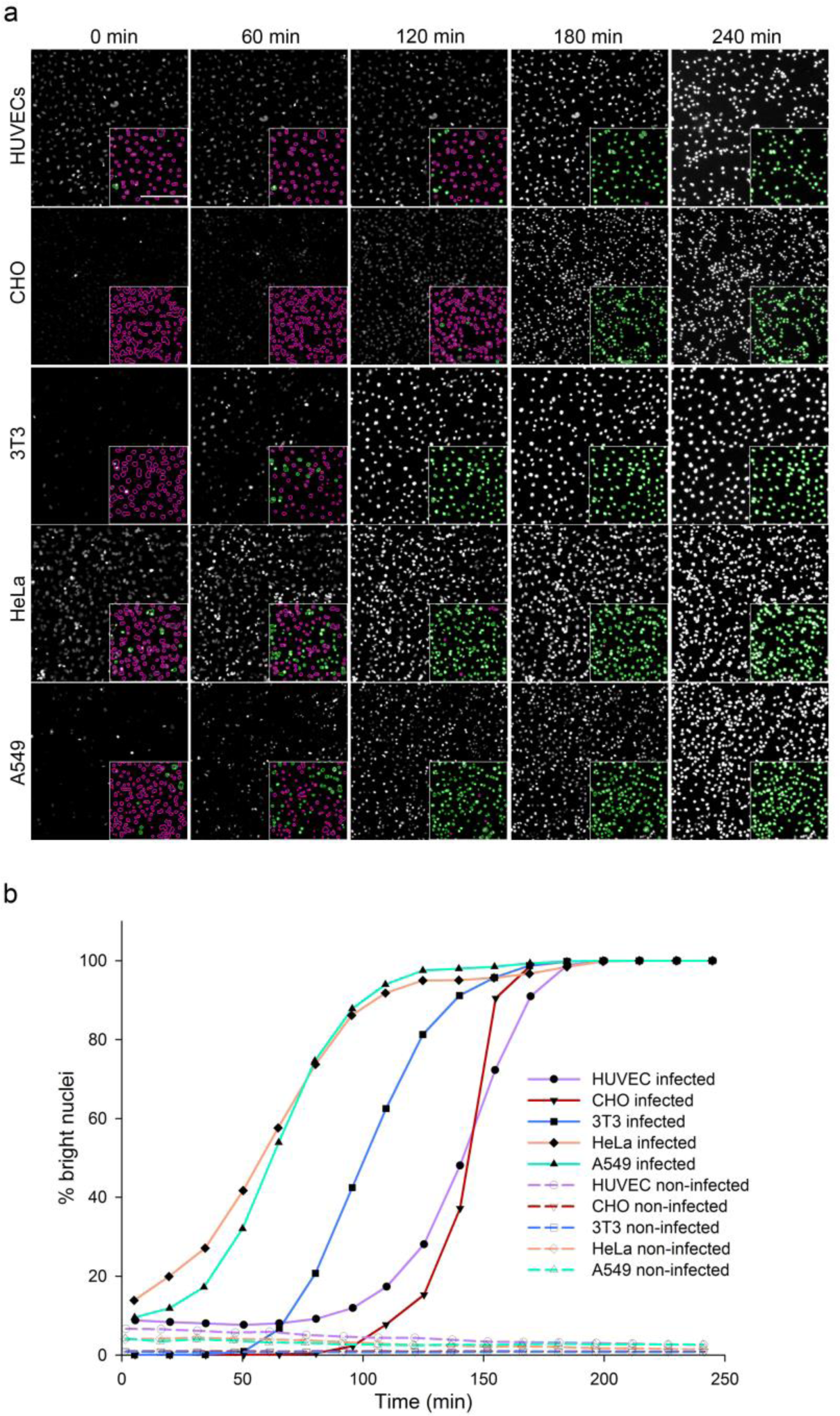
Live-imaging quantification of *P. aeruginosa* cell infection by fluorescence intensity measurement of Hoechst-labelled nuclei. HUVEC, CHO, 3T3, HeLa and A549 cells were labelled with vital-Hoechst prior to infection with *P. aeruginosa* and monitored by microscopy at a 5x magnification. a) Cell nuclei observed by fluorescent labelling and automatically segmented (insert). Nuclei with intensities below a fixed threshold were delineated in magenta while those above the threshold were delineated in green. The scale bar shown on the first image of the HUVECs cells corresponds to 200 μm. b) Kinetic plots representing the percentage of nuclei with intensities above the thresholds in the images taken every 15 min.

### Detection of bright nuclei is highly discriminant

We then focused our analysis on HUVECs which are particularly relevant because they are primary human cells, forming a polarized monolayer. The action of *P. aeruginosa* on HUVECs have been extensively studied by cellular biology and the morphological changes observed during infection have been described by microscopy approaches^9,11–15^. Furthermore, a “cell area” assay based on the quantification of fixed cells’ area by immunofluorescence staining was previously reported^9,14^. Therefore, we investigated how the CLIQ-BID method compares to the previously published method which is based on immunofluorescence cell staining. For this purpose, HUVECs were infected for different time periods, their nuclei were observed with vital Hoechst staining and then immediately fixed, immunostained and observed at the same position in the wells. A comparison of the results obtained with the CLIQ-BID and “cell area” quantification methods is presented in Figure 3. Nuclei were segmented and discriminated based on their fluorescence intensities (Fig. 3 a, Hoechst and bright nuclei images) while the cells’ area was quantified based on thresholding of the vinculin staining (Fig. 3a, Vinculin and cell area images). Indeed, the area covered by the cells decreased during infection with *P. aeruginosa*, in agreement with previous observations^14^. Furthermore, the plots of the percentage of bright nuclei and the percentage of the field area cleared by the cells after different infection durations exhibit a similar trend and the correlation coefficient between the readouts of the image pairs was 0.96. However, the standard deviations were much lower for the percentage of bright nuclei than for the cell area (Figure 3b).

**Figure 3:**
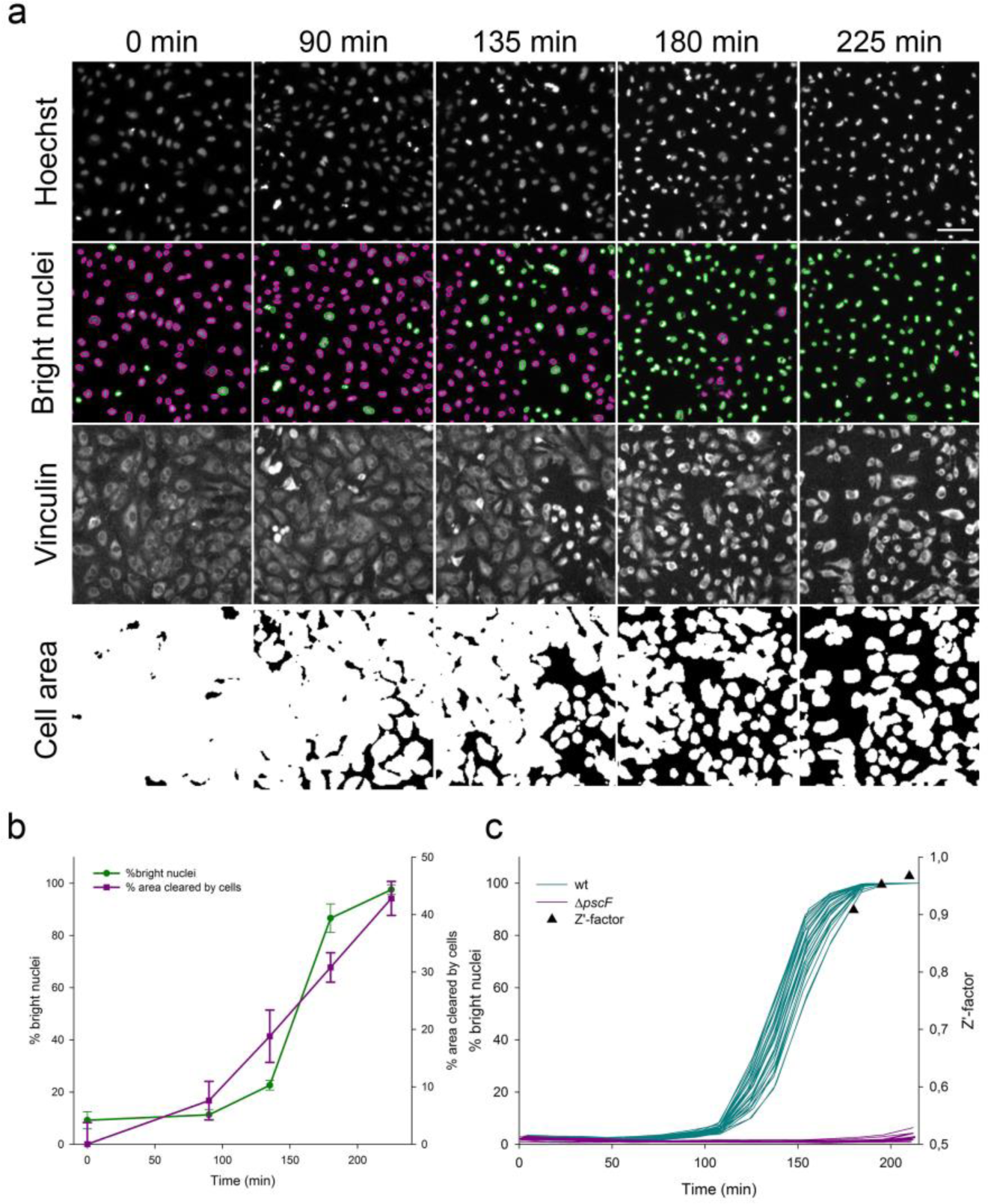
Comparison of the CLIQ-BID and “cell area” methods. HUVECs were infected with *P. aeruginosa* for different durations; images of the nuclei were acquired before cell fixation, immunostaining and acquisition of cell area images. a) Image sets at different time points for i) Hoechst: cell nuclei; ii) Bright nuclei: nuclei automatically segmented and sorted for intensities below (red) or above (green) a fixed threshold; iii) Vinculin: cell area detected with a cytoplasmic vinculin staining; iv) Cell area: automated thresholding of the immunostaining allowing the calculation of the field area covered by the cells. The scale bar shown on the last nuclei image corresponds to 50 μm. b) Plots of the percentage of nuclei with intensities above the thresholds and of the percentage of area cleared by the cells after different infection durations. Error bars represent the standard deviation (n=8).

In order to confirm that the CLIQ-BID method reduces the variation between individual data points, the comparison experiment was repeated with two modifications: i) the number of replicate wells was set to 30 per condition and ii) the HUVECs were infected for 3 hours either with a wild-type *P. aeruginosa* strain or the Δ*pscF* strain, which is unable to produce the T3SS toxin injection needle and is therefore deficient for a functional T3SS. This experiment substantiated that the CLIQ-BID method produces more robust and discriminant results with a Z’-factor equal to 0.89 versus 0.14 for the cell area method (Supplementary Table S2).

The high Z’-factor obtained with the CLIQ-BID method indicated that it is well-suited for high throughput screening. To further examine this possibility, cell infections with wild-type or Δ*pscF P. aeruginosa* strains were compared in 96- and 384-well plates. The live-imaging monitoring of nuclei intensities allowed the obtaining of reproducible kinetics curves in the 48 replicates (Supplementary Fig. S2). Indeed, the Z’-factor values obtained by comparing the wild-type and the T3SS deficient strains were close to or above 0.9 for both plate formats. Importantly, an additional 384-well plate was inoculated with overnight *P. aeruginosa* cultures, as opposed to exponential phase cultures, which is currently used to detect T3SS activity (Supplementary Fig. S2). The removal of the subculture step significantly reduces the handling procedure in the perspective of bacteria or molecule library screens. Despite higher variation than with exponential cultures, the Z’-factor displayed values close to 0.8, higher than the gold-standard of 0.7 above which a library could be screened in a single replicate with an acceptable risk of false-positives and -negatives. Taken together, these results indicate that the newly-developed CLIQ-BID method is adapted for HTS strategies.

In the search for a global descriptor of each kinetics plot, the Area Under the Curve (AUC) was selected. As expected, large differences were observed between AUC obtained from wells infected with wild-type or T3SS deficient strains (Supplementary Fig. S3). Indeed, the statistical analysis showed that the AUC can robustly discriminate between infections by these two strains, with Z’-factor values of 0.77 and 0.63 in 96- and 384-well plates, respectively (Supplementary Table S3).

### Bright nuclei detection enables potent screening strategies

Considering the encouraging results, the potential of the method to screen for inhibitors of bacteria-induced cell damage was further investigated on a panel of molecules. In this proof-of-concept experiment, compounds targeting the bacteria or the eukaryotic cells were tested, along with siRNA. A major improvement was also made with the use of bacteria constitutively expressing GFP in their cytosol. Measuring the global GFP fluorescence increase in the wells thus enables the detection of possible bacteriostatic/bactericidal properties of the tested compounds. These experiments were therefore analysed by plotting the kinetics curves of bright nuclei (cell toxicity) and GFP fluorescence (bacterial growth), as represented in Figure 4.

**Figure 4:**
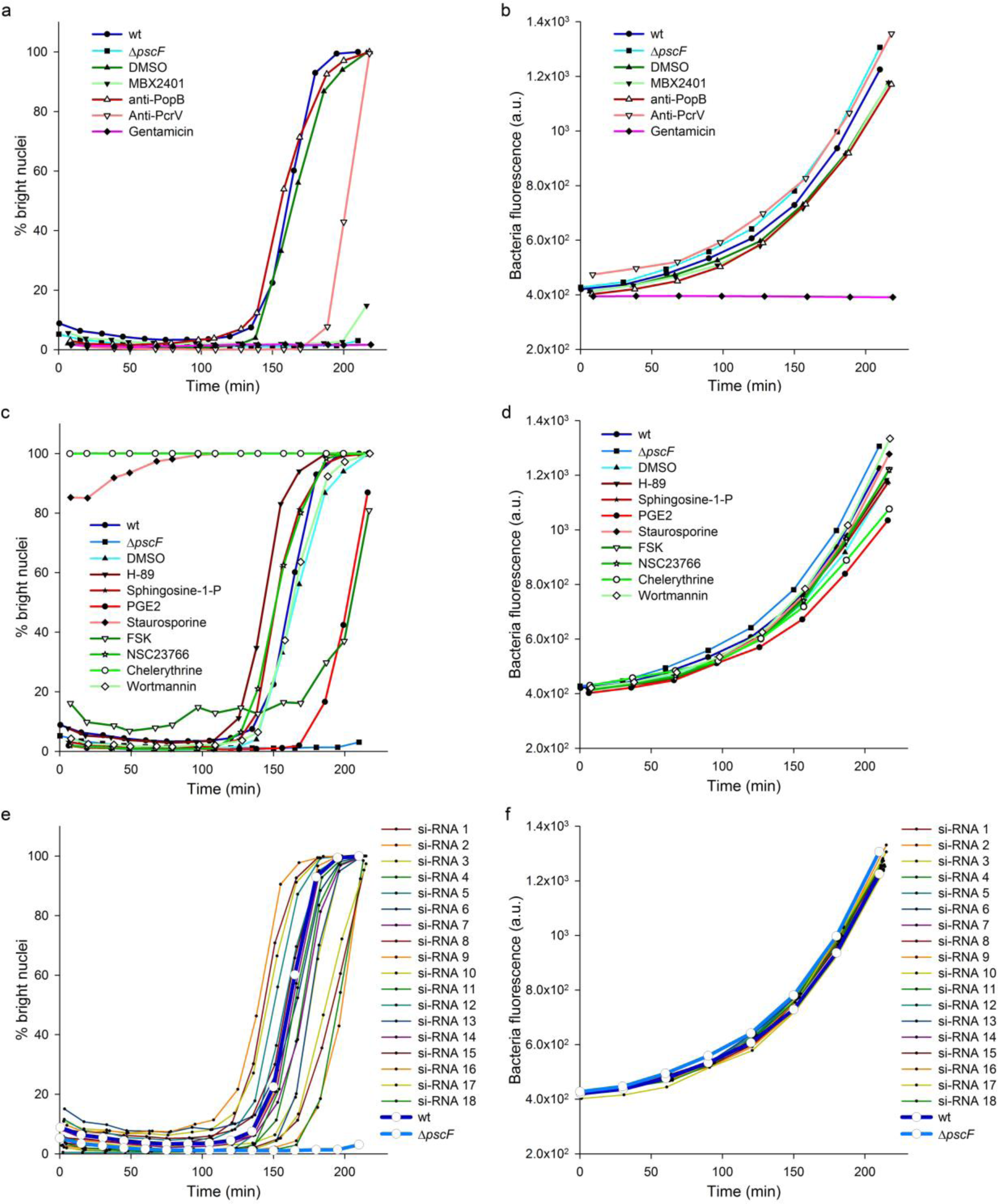
Screening of a panel of molecules. HUVECs were infected with *P. aeruginosa* wild-type strain in the presence of molecules targeting the bacteria (a, b) or targeting the eukaryotic cells (c, d) and their respective controls. To assess the effect of siRNA transfection in the cells, HUVECs were transfected two days before infection (e, f). A strain deficient for the production of the T3SS needle subunit (Δ*pscF*) was used as control. The kinetics of bright nuclei appearance (a, c, e) and bacteria growth (b, d, f) were simultaneously recorded by live-imaging and analysed. a.u. = arbitrary units.

Cell damage were significantly delayed by known inhibitors of *P. aeruginosa* T3SS, like the small molecule MBX2401 or polyclonal antibodies raised against the tip protein PcrV^16,17^. Conversely, neither DMSO nor antibodies targeting the translocator PopB had any effect on the cell infection, as expected (Fig. 4a). Among molecules targeting the eukaryotic cells, the prostaglandin PGE2 and forskolin, a cAMP inducer recently shown to inhibit *P. aeruginosa* T3SS effects through Rap1 activation^12^, significantly delayed the kinetics (Fig. 4c). On the other hand staurosporine and chelerythrine, two potent cytotoxic inhibitors of protein kinases^18,19^, immediately induced the appearance of bright nuclei. Other compounds moderately accelerated cell damage during infection, namely H-89 (protein kinase A inhibitor), sphingosine-1-P (signalling phospholipid) and NSC23766 (Rac1 inhibitor), while wortmannin (PI3-K inhibitor) had no effect. Of note, cells incubated with H-89, sphingosine-1-P and Forskolin in the absence of bacteria exhibited a higher basal level of bright nuclei (Supplementary Fig. S4). This moderate toxicity could explain the accelerated kinetics observed when cell were infected in the presence of H-89 and sphingosine-1-P and the relatively high baseline observed with foskolin at the beginning of cell infection. Finally, cells were grown after transfection with arbitrarily chosen siRNAs from a laboratory collection and were subsequently infected. From the 18 tested siRNAs, some had no effect while some exhibited promoting or inhibiting activities (Fig. 4e). The validation and the biological investigation of the role of their targets are beyond the scope of this work.

Observing the effect of a particular treatment on bacteria growth (Fig. 4b, d, f) allowed the determination as to whether any virulence inhibition was related to an antibiotic effect. Indeed, none of the eukaryotic nor T3SS-specific targeting compounds exhibited a bacteriostatic effect, while the antibiotic gentamicin prevented bacterial growth and, consequently, host cell intoxication. Furthermore, the synthetic descriptors of the cell toxicity and bacterial kinetics curves (respective AUCs) permitted the straightforward statistical comparison of the kinetics by one-way ANOVA (Supplementary Fig. S5). In conclusion, the developed method enables the identification of promotional or inhibitory effects from a variety of molecular categories (small organic molecules, antibodies and siRNAs) and allows one to simultaneously counter-screen for antibiotic effects.

Finally, the method was employed to compare the effects of different bacterial strains and bacterial species. For this purpose, HUVECs were infected with 16 different bacteria and the intensities of cell nuclei were monitored by live-imaging. Kinetics plots and the corresponding AUCs (Fig. 5) show the diverse virulence potential of these bacteria. Among the *P. aeruginosa* strains, the PP34 strain, injecting through its T3SS the powerful phospholipase toxin ExoU^3,7,20^, was the most active followed by IHMA87 and PAO1Δ *pscD*⸬*exlBA* (PAO1 *exlBA*) strains expressing the recently discovered Two Partner Secreted toxin ExlA^21–23^ and the CHA reference strain used throughout this study. Of interest, the *Serratia marcescens* strain expressing the ShlA toxin^24^ homolog to ExlA was as active as the PP34 strain, while the isogenic mutant strain that has a transposon inserted into the *shlB* gene^25^, did not induce the appearance of bright nuclei. In *Staphylococcus aureus,* the 8325-4 strain exhibited lower effects than the closely related USA300 BEZIER and SF8300 strains, both from the USA300 lineage known to be highly virulent^26^. This workflow was also successful in detecting cellular damage caused by *Yersinia enterocolitica*, which correlated with its T3SS since the mutant depleted of T3SS toxins exhibited a significantly lower activity. Finally, five bacteria species did not display measurable effects toward the eukaryotic cells under the used experimental conditions, notably the relatively short time span of infection and low MOI. Of interest, the statistical analysis of AUCs derived from the kinetics plots of each replicate confirmed the depicted intra-species differences (Fig. 5b). The sigmoid curves obtained with the virulent bacteria display different shapes, notably regarding the lag phase and the slope. These parameters are not expected to be correlated because different virulence mechanisms triggers effects with different delays and different degrees of synchronicity in the target cell population. The AUCs calculation does not give access to these variations and it is conceivable that they could compensate, resulting in some cases in similar AUCs for different curve shapes. Therefore, the curves of each replicate of the same experiment were fitted using a sigmoid equation and the calculated inflection points and curve steepness were represented on a XY plot for each bacteria species (Fig. 5c). This analysis clearly highlights inter- and intra-species differences while similar strains within *P. aeruginosa* (PAO1 *exlBA* and IHMA87) and *S. aureus* (SF8300 and USA300) species clustered together. Overall, real-time imaging allowed the observation of different cell-damage kinetics, which are in agreement with what is expected for the corresponding bacteria featuring diverse toxins and virulence mechanisms.

**Figure 5:**
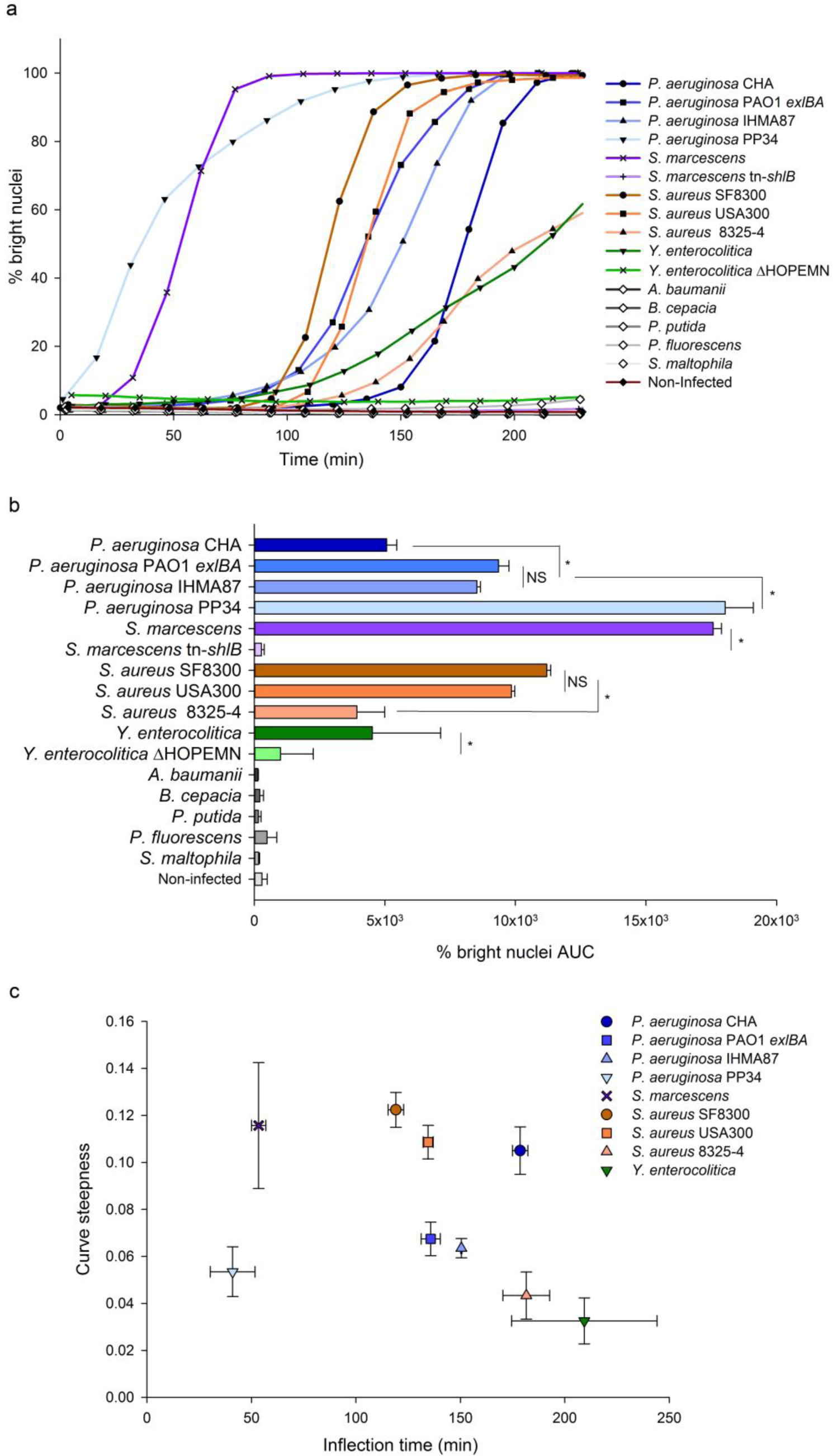
Comparison of cell damage kinetics with different bacteria. HUVECs were infected with 16 different bacteria and monitored by live-imaging. The percentages of bright nuclei were used to derive kinetics plots (a) and the corresponding Area Under the Curves (AUC) (b). Each point in the kinetics plots correspond to the means of triplicates and AUC histograms represent the means of the AUCs obtained for the kinetics of each replicate. Error bars represent the standard deviation (n=3). Stars indicate statistically significant differences between strains of the same species and NS a non-significant difference (one-way ANOVA, P < 0.05). Each of Tthe kinetics plot replicatess obtained with different bacterial species inducing “bright nuclei” were fitted with sigmoid model curves and the inflection point and curve steepness were calculated and represented as XY pairs (c). Means and standard deviations are represented.

## DISCUSSION

The new high-throughput image analysis strategy described in this work exhibits great potential for monitoring cell damage induced by bacteria, as well as by other mechanisms. Through nuclei monitoring, the method allows the observation of cell-reaction to Grampositive and Gram-negative bacteria secreting or injecting toxins. Furthermore, both cell lysis induced by plasma-membrane targeting toxins ExoU and ExlA^20,22^, as well as cell shrinkage induced by ExoS^9,27^ were readily detected. Of note, this method can also be employed to reveal cytotoxicity or cell stress from a variety of origins, as exemplified by the detection of the effects of chelerythrine and staurosporine known to promote apoptosis. It is thus able to detect cell shrinkage, necrosis and apoptosis.

The developed method allows cost-effective kinetic measurements of bacteria-induced action on several cell lines, requiring a simple staining procedure, a microscope and an image-analysis software. Detection of nuclei and quantification relies on the widely-used Hoechst 33342, an inexpensive vital stain of cell nuclei used for almost four decades^28^. After image acquisition on the microscope, images may be analysed with standard software like Cell Profiler, ImageJ or Fiji^29–31^, using basic algorithms to delineate nuclei and measure their fluorescence intensities. In our work, we used an automated microscope along with its proprietary analysis software to demonstrate the great potential of this method for High Content Screening.

Indeed the Z’-factor values obtained in different assay configurations were often above 0.8. This statistical descriptor reflects the quality of a screening method, i.e. its ability to identify “hits”^10^. Screening of libraries are undertaken only if the Z’-factor is above 0.6 and a value above 0.7 is considered to be fully satisfactory. The elevated Z’-factor value obtained in 384- well microplates with fresh or overnight cultures of bacteria indicated that this assay could be employed to screen large bacterial mutant or eukaryotic CRISPR/Cas9 libraries. Furthermore, the screening approach was successfully applied to a set of antibodies and small molecules targeting either bacteria or eukaryotic cells and to a panel of siRNAs. Indeed, it identified three inhibitory activities among the tested small molecules and antibodies: the MBX2401 drug and anti-PcrV antibodies, both known to inhibit *P. aeruginosa* T3SS^16,17^, and forskolin, known to counteract the T3SS effect in eukaryotic cells^12^. This represents a proof of concept for a screening strategy. Moreover, the morphological readout of the method (brightness of the nuclei) is a downstream event in the infection process, enabling the screening of molecules that could target early or late events in the bacteria or the host.

In addition, the simultaneous quantification of GFP-expressing bacteria allows one to counter-screen, in the same test, for bacteriostatic/bactericidal activities. Moreover, the same CLIQ-BID method can be used in the absence of bacteria to detect the potential deleterious effects of screened compounds. Therefore, this method can be the basis of a powerful 3-in-1 approach to rapidly identify treatments inhibiting bacteria virulence without affecting their growth capacities or the eukaryotic cells’ integrity. This is of particular interest since the search for antivirulence treatments is receiving growing attention because they are thought to reduce the risk of resistance emergence and microbiome destabilization^32,33^.

The increasing accessibility to HCS/HCA equipment, with the help of the simple and cost-effective CLIQ-BID method described here, should foster the understanding of bacterial virulence as well as other scientific areas where early cell damage and cell stress are of interest.

## METHODS

### Bacteria strains

*Pseudomonas aeruginosa* strains CHA, PAO1 Δ*pscD*⸬*exlBA*, IHMA87 and PP34; and *Serratia marcescens* Db11 and Db11-tn-*shlB* were from our lab collection and previously published^14,21,22,25^. *Staphylococcus aureus* strains SF8300, USA300 BEZIER and 8325-4 were a kind gift from Dr Karen Moreau. Other strains were *Yersinia enterocolitica* E40 and ΔHOPEMN^17^, *Acinetobacter sp. genomospecies 13* ATCC 23220, *Burkholderia cepacia* ATCC 17616, *Pseudomonas putida* KT2442, *Pseudomonas fluorescens* BG1 (environmental isolate, gift from Dr John Willison) and *Stenotrophomonas maltophila* (ATCC 13637). Bacteria were grown in LB (Luria Bertani – Difco) except for *S. aureus* which were grown in BHI (Brain Heart Infusion - Difco). *Y. enterocolitica* strains were grown at 28 °C and the other strains at 37 °C. Unless otherwise stated, cultures were grown overnight under shaking at 300 rpm and then diluted in fresh media to an optical density measured at 600 nm (OD_600_) of 0.1. When cultures reached OD_600_ of 1, typically after 2.5 h of growth under shaking, bacteria were mixed with eukaryotic cells at a multiplicity of infection (MOI) of 10.

### Chemicals and antibodies

Hoechst 33342, Gentamicin, H-89, Sphingosine-1-P, Prostaglandin E2, Staurosporine, Forskolin, NSC23766 and Wortmannin were from Sigma-Aldrich and chelerythrine from Merck Millipore. MBX2401, an inhibitor of the Type Three Secretion System (T3SS) from *P. aeruginosa*^16^ was synthesized as previously described^34^. Antibodies raised against *P. aeruginosa* PopB and PcrV (anti-PopB and anti-PcrV) were obtained in our laboratory and previously characterized^17^. Antibodies to Vinculin were from Santa Cruz and the secondary antibodies coupled to Alexa 488 were purchased from Molecular Probes.

### Cell culture

Human umbilical vein endothelial cells (HUVECs) were isolated according to previously described protocols^9^. The use of umbilical cords for scientific purposes is authorized by the L1211-2 act from the French Public Health Code. Written informed consent was obtained from each woman who donated an umbilical cord. The privacy of the donor’s personal health information was protected. Recovered cells were cultured in endothelial-basal medium 2 (EBM-2; Lonza) supplemented as recommended by the manufacturer. A549 (CCL-185) and HeLa (CCL-2) cells were grown in RPMI supplemented with 10% foetal calf serum. 3T3 (CRL-2752) and CHO K1 (CCL-61) cells were grown in DMEM and F12 medium, respectively, supplemented with foetal calf serum.

### siRNA

Cells were seeded at 12,500 cells per/well in a 96-well plate and transfected with siRNAs using Lipofectamine™ RNAiMax transfection reagent (Thermo Fisher Scientific), according to the Reverse Transfection manufacturer’s protocol. Briefly, 2.5 pmol of siRNA were diluted in 10 μl of supplemented EBM-2 and mixed with 0.2 μl of transfection reagent, pre-diluted in 9.8 μl of supplemented EBM-2. After 15min at room temperature, the complexes were added to the cells in a final volume of 100 μl of supplemented EBM-2. Cells were used 48 h later.

### Cell treatments, Hoechst staining and infection

Black μclear 96-well plates (Greiner) were seeded at 12,500 cells per/well. Black μclear 384- well plates (Greiner) were seeded at 3,000 cells per well. Cells were used 48 h later to obtain highly confluent monolayers. Medium was replaced 3 h before infection with fresh medium containing Hoechst 33342 (1 μg/ml). After incubation during 1 h, two washes with 80 μl of fresh non-supplemented EBM-2 medium. All media were pre-heated at 37 °C.

For pharmacological and antibody treatments, medium was replaced 30 min before infection with 80 μl of fresh medium supplemented with: gentamicin 200 μg/ml, H-89 10 μM, sphingosine-1-P 2 μg/ml, prostaglandin E2 1 nM, staurosporine 1 μM, forskolin 10 μM, NSC23766 50 μM, wortmannin 1 μM, chelerythrine 1 μM, MBX2401 and MBX2402 30 μM. When applicable, the final DMSO concentration was 0.5%. Sera containing antibodies directed against PopB and PcrV were diluted to a final concentration of 5%.

Cells were infected at a multiplicity of infection (MOI) of 10 by adding 20 μl of EBM-2 containing a 5x concentrated bacteria suspension. Plates were immediately observed by Arrayscan microscopy.

### Automated High Content Imaging and High Content Analysis (HCA)

The image acquisitions were performed on an automated microscope ArrayScanVTI (Thermo Scientific) using a Zeiss 20x (NA 0.4) LD Plan-Neofluor or a Zeiss 5x (NA 0.25) Fluar air objectives. In 96-well plates, four images per well were acquired with the 20x or 5x objectives and one image per well was acquired with the 5x objective in 384-well plates. The dichroic mirror used for Hoechst staining was BGRFR-386/23 nm and BGRFR-brightfield for transmitted light imaging. Exposure times were set to reach 40% of intensity saturation in the reference wells at the beginning of the experiment. The microplate was maintained at 37°C and 5% CO2 in the ArrayScan Live Cell Module and images were automatically acquired every 15 min for up to five hours. Indicated times refer to the actual time of image acquisition and are adjusted for delay between wells due to the plate displacement. Typically five minutes are required to scan a complete 96-well plate.

Quantification of nuclei parameters was made using the Cell Health Profiling Bio-Application of Thermo Scientific HCS Studio v6.5.0. Each nucleus was detected in the Hoechst channel with the isodata thresholding method. Border-touch nuclei were rejected from each image. Nuclei area and nuclei average intensity features, respectively named ObjectAreaCh1 and ObjectAvgIntenCh1, were automatically calculated. When indicated in the text, a threshold was applied on the ObjectAvgIntenCh1 feature to discriminate the population of cells with bright nuclei. This threshold was set to 2,100 or 1,200 fluorescence arbitrary units (a.u.) for 20x or 5x magnification images respectively, and the proportion of cells with bright nuclei were automatically calculated and named %HIGH_ObjectAvgIntenCh1. The arbitrary units correspond to the raw fluorescence intensities obtained through the HCS Studio software.

### Immunofluorescence staining and quantification

Cells were washed, fixed with 4% paraformaldehyde for 15 min, permeabilized with 0.5% Triton X-100 for 5 min and labelled with a mouse anti-Vinculin 7F9 primary antibody (Santa Cruz) and donkey anti-mouse Cy3 secondary antibody (Jackson ImmunoResearch Laboratories) for 1 h each. Nuclei were labelled with Hoechst 33258 (10 μg/ml, Sigma-Aldrich). Under these conditions, total cellular vinculin is detected yielding a whole-cell labelling. Images were captured using an automated microscope ArrayScanVTI with the 5x magnification objective, in the BGRFR-549/15 nm channel, and treated with ImageJ software. Briefly, images of vinculin staining were binarized and the total cell area was calculated for each image. To facilitate interpretation, results were shown as the percentage of the field area cleared by the cells.

### Statistics

To evaluate the quality of the assay and its ability to identify “hits”, the Z’-factor was calculated using the following equation, as described by Zhang et al ^10^:

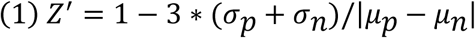

where *σ_p_* and *σ_n_* are the standard deviations of the positive and negative conditions, respectively, and *μ_p_* and *μ_n_* are the means of the positive and negative conditions, respectively.

Statistical analyses were performed using SigmaPlot 12.5 (Systat software) for the comparison of multiple groups by one-way ANOVA (two-tailed). When appropriate, *post hoc* tests were Tukey or Dunnett for multiple comparisons or comparison to the control group, respectively.

In figure legends, n represents the number of well replicates.

Curve fitting was done with SigmaPlot 12.5 using the following equation:

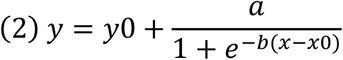

where *y0* and *a* are the minimal and the range values of the bright nuclei percentage, respectively, and *x0* and *b* respectively correspond to the inflection time and the curve steepness.

## Data availability

The datasets generated during and/or analysed during the current study are available from the corresponding author on reasonable request.

## ACKNOWLEDGMENTS

Y.W. was a recipient of a postdoctoral fellowship from the Fondation pour la Recherche Médicale. This project was in part supported by grants from the AVIESAN T3SS (ANR PRP1.4), the Laboratory of Excellence GRAL (ANR-10-LABX-49-01) and the Agence Nationale de Recherche (ANR-15-CE11-0018-01). We acknowledge the Labex GRAL and IBiSA for financial support to the CMBA platform. Authors are grateful to Charles Ross Dunlop for proofreading the manuscript.

## AUTHORS CONTRIBUTIONS

Y.W. and E.F. designed experiments, Y.W., S.B., P.H. and E.F. performed experiments, I.A. and E.S. contributed reagents and methods, Y.W., E.S., P.H., I.A. and E.F. analyzed and discussed the data and E.F. wrote the manuscript. All authors contributed to and edited the manuscript.

## ADDITIONAL INFORMATION

Competing Interests: The authors declare that they have no competing interests.

